# Rac1 and Nectin3 are essential for PCP-directed axon guidance in the peripheral auditory system

**DOI:** 10.1101/2024.06.05.597585

**Authors:** Shaylyn Clancy, Nicholas Xie, Tess Eluvathingal Muttikkal, Jasmine Wang, Esha Fateh, Margaret Smith, Phillip Wilson, Matthew Smith, Arielle Hogan, Ann Sutherland, Xiaowei Lu

## Abstract

Our sense of hearing is critically dependent on the spiral ganglion neurons (SGNs) that connect the sound receptors in the organ of Corti (OC) to the cochlear nuclei of the hindbrain. Type I SGNs innervate inner hair cells (IHCs) to transmit sound signals, while type II SGNs (SGNIIs) innervate outer hair cells (OHCs) to detect moderate-to-intense sound. During development, SGNII afferents make a characteristic 90-degree turn toward the base of the cochlea and innervate multiple OHCs. It has been shown that the Planar Cell Polarity (PCP) pathway acts non-autonomously to mediate environmental cues in the cochlear epithelium for SGNII afferent turning towards the base. However, the underlying mechanisms are unknown. Here, we present evidence that PCP signaling regulates multiple downstream effectors to influence cell adhesion and the cytoskeleton in cochlear supporting cells (SCs), which serve as intermediate targets of SGNII afferents. We show that the core PCP gene Vangl2 regulates the localization of the small GTPase Rac1 and the cell adhesion molecule Nectin3 at SC-SC junctions through which SGNII afferents travel. Through *in vivo* genetic analysis, we also show that loss of Rac1 or Nectin3 partially phenocopied SGNII peripheral afferent turning defects in *Vangl2* mutants, and that Rac1 plays a non-autonomous role in this process in part by regulating PCP protein localization at the SC-SC junctions. Additionally, epistasis analysis indicates that Nectin3 and Rac1 likely act in the same genetic pathway to control SGNII afferent turning. Together, these experiments identify Nectin3 and Rac1 as novel regulators of PCP-directed SGNII axon guidance in the cochlea.

**Significance statement:** Planar Cell Polarity (PCP) signaling plays a non-autonomous role in the guidance of type II spiral ganglion neuron (SGNII) afferent projections that innervate cochlear hair cells. However, little is known about the underlying mechanisms. Here, we identify the small GTPase Rac1 and the cell adhesion molecule Nectin3 as two downstream effectors of PCP signaling in SGNII afferent guidance. We show that PCP signaling regulates Rac1 and Nectin3 localization in cochlear supporting cells that serve as intermediate targets for SGNII afferents and that Rac1 and Nectin3 likely act in the same genetic pathway to non-autonomously regulate SGNII afferent guidance. These findings significantly advance our understanding of auditory circuit assembly and shed light on PCP-directed axon guidance mechanisms.

## Introduction

The mammalian auditory sensory epithelium, the Organ of Corti (OC), is a stratified epithelium consisting of sensory hair cells (HCs) and multiple types of supporting cells (SCs) whose cell bodies are positioned below those of the HCs. In the OC, HCs are interdigitated by the phalangeal processes of neighboring SCs, forming a highly organized cellular mosaic **(Fig. 1A, B)**. Atop each HC sits a V-shaped stereociliary hair bundle (HB). Mechanical deflection of the HB in response to sound depolarizes the HC, converting sound information into electrical signals that are transmitted via the spiral ganglion neurons (SGNs). SGNs are bipolar afferent neurons with a peripheral axon that innervates HCs and a central axon that projects to the auditory brainstem. Type I SGNs transmit primary sound information from the inner HCs (IHCs); type II SGNs (SGNIIs) innervate the three rows of outer HCs (OHCs) and are thought to play a role in detecting moderate-to-intense sound and volume processing (Zhang and Coate, 2017; Weisz et al., 2021). While multiple type I SGN afferents innervate a single IHC, the SGNII peripheral afferent projections travel through SCs before they make striking 90-degree turns toward the base of the cochlea while extending apically and forming *en passant* synapses with multiple OHCs.

**Figure 1:**
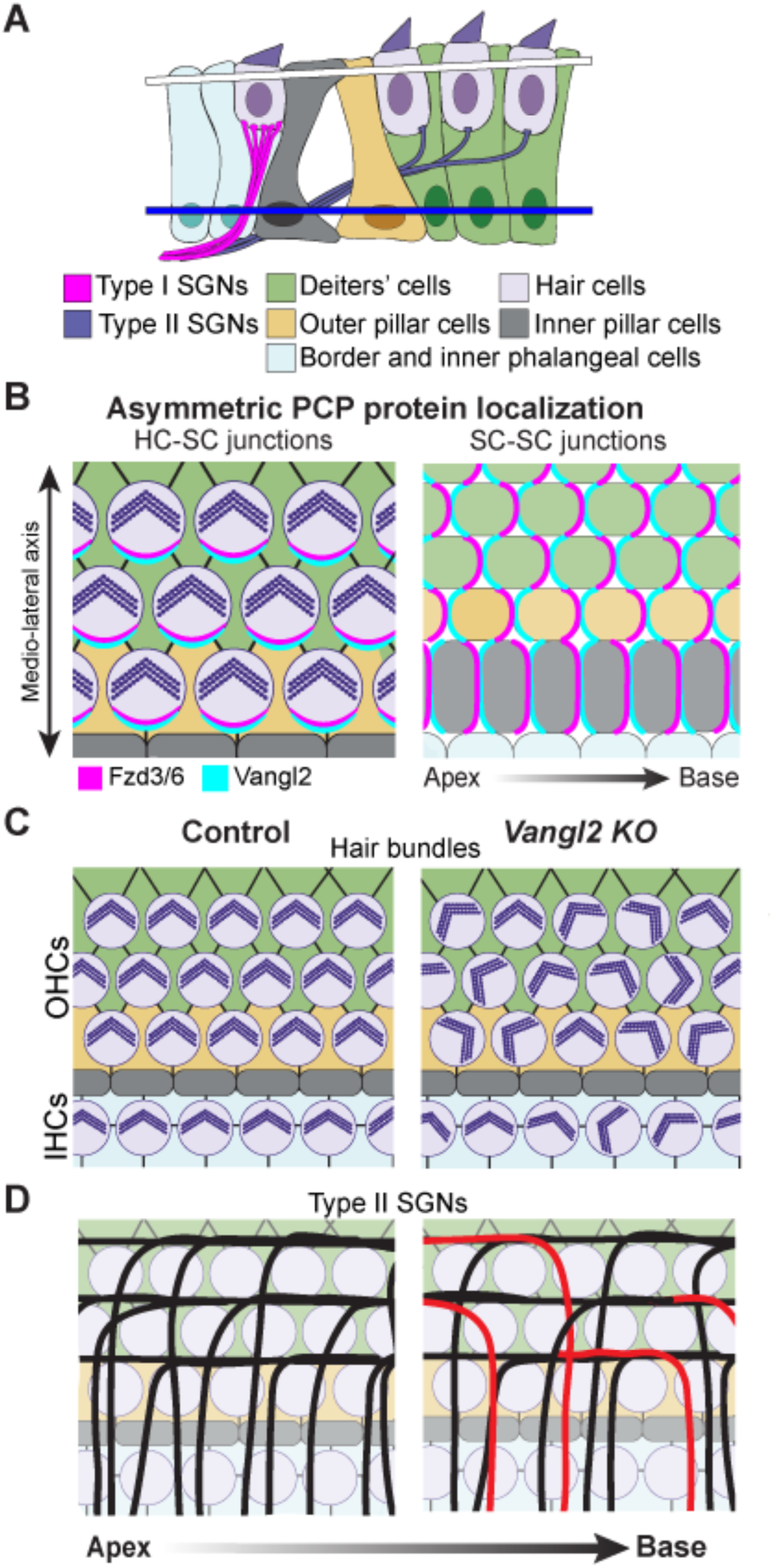
PCP signaling in the cochlea. **A**, A cross-sectional view of the cochlea demonstrating Type I and II SGN afferent innervation patterns. The white bar indicates the HC-SC junctional level, while the blue bar indicates the SC-SC junctional level. **B,** Schematic diagrams showing asymmetric localization of Fzd3/6 and Vangl2 at the HC-SC and SC-SC junctional levels. **C, D**, Loss of PCP signaling leads to misoriented HBs (**C**) and a percentage of SGNII afferents to turn errantly toward the cochlear apex (shown in red) (**D**).

The evolutionarily conserved planar cell polarity (PCP) pathway aligns cell polarity across a tissue plane and controls polarized cell behaviors during vertebrate tissue morphogenesis (Butler and Wallingford, 2017). Critical for their function in aligning planar polarity, PCP proteins form two asymmetric protein complexes that serve as molecular “Velcro” across intercellular junctions. In the inner ear, it has been shown that intercellular PCP signaling in the cochlear epithelium aligns HB orientation and guides SGNII afferent turning (Tarchini and Lu, 2019; Deans, 2022), mediated by asymmetric PCP protein complexes at HC-SC and SC-SC junctions, respectively (**Fig. 1B-D)**. At HC-SC junctions **(Fig. 1A, white bar),** PCP proteins such as Van Gogh-like 2 (Vangl2) and Frizzled (Fzd) 3 and 6 are asymmetrically localized along the medial-lateral axis of the cochlear duct to align HB orientation (Montcouquiol et al., 2003; Wang et al., 2006) **(Fig. 1B, C)**. Interestingly, at SC-SC junctions **(Fig. 1A, blue bar)**, Vangl2 and Fzd3/6 are asymmetrically localized on the apex- and base-facing junctions, perpendicular to the medial-lateral axis, to steer the SGNII afferents toward the cochlear base (Ghimire et al., 2018; Ghimire and Deans, 2019) **(Fig. 1B, D)**. However, the underlying mechanisms for SGNII afferent guidance are currently not well understood. In other systems, PCP signaling has been shown to mediate tissue morphogenesis through regulation of the cytoskeleton and cell adhesion (Butler and Wallingford, 2017; Dreyer et al., 2022). Thus, we wanted to investigate whether PCP signaling is acting through cytoskeletal and cell adhesion regulators in the OC to influence SGNII afferent guidance in a non-cell-autonomous manner.

We reasoned that molecules known to be expressed in SCs and previously implicated in regulating HB orientation likely also play a role in influencing SGNII afferent turning. Our lab has previously shown the small GTPase Rac1, a central regulator of cytoskeletal remodeling, to be necessary for proper HB orientation and morphology (Grimsley-Myers et al., 2009). In cultured cells, Vangl2 interacts with and localizes Rac1 at adherens junctions without altering Rac1 activity (Lindqvist et al., 2010). Given these data, we hypothesized that Rac1 is a downstream effector of Vangl2 for SGNII afferent guidance.

Similarly, Nectin3, a member of the immunoglobulin superfamily of cell adhesion molecules (IgCAMs) expressed in SCs, has been shown to form heterophilic adhesion at HC-SC junctions with Nectin1 expressed in HCs (Togashi et al., 2011). Nectin-mediated cell adhesion is required for establishing the characteristic checkerboard layout of the OC, as well as HB orientation and morphology (Togashi et al., 2011; Fukuda et al., 2014). Therefore, Nectin3 may participate in PCP signaling and play a role in SGNII afferent guidance.

Using immunolocalization and genetic loss and gain of function approaches, we show that Rac1 and Nectin3 are localized to SC-SC contacts and required for SGNII afferent turning *in vivo* and likely function in the same pathway in SCs to guide SGNII afferents. These results reveal a Vangl2-Nectin3-Rac1 feedback regulatory circuit that presents guidance cues for SGNII afferent turning in the murine cochlea.

## Results

### Rac1 is localized to SC-SC junctions and its localization is regulated by Vangl2

To determine a potential non-autonomous role of Rac1 in guiding SGNII afferents, we began by immunofluorescence (IF) labeling of total Rac1 protein at the SC-SC junctional level. In the control OC at embryonic day (E) 18.5, Rac1 was localized at the SC-SC contacts, with a modest but significant increase in Rac1 IF intensity in the middle and apical turns compared to the cochlear base **(Fig. 2A, B)**. Interestingly, in *Vangl2* knockout (KO, *Vangl2^KO^*) cochleae, there was a significant increase in Rac1 IF intensity at the SC-SC junctional level along the entire length of the cochlea **(Fig. 2A, B)**. In contrast to the control, there was no significant tonotopic difference in Rac1 IF intensity **(Fig. 2B)**. Thus, altered Rac1 localization at SC-SC junctions in *Vangl2^KO^* cochleae suggests that Rac1 is a potential PCP effector in guiding SGNII afferents.

**Figure 2:**
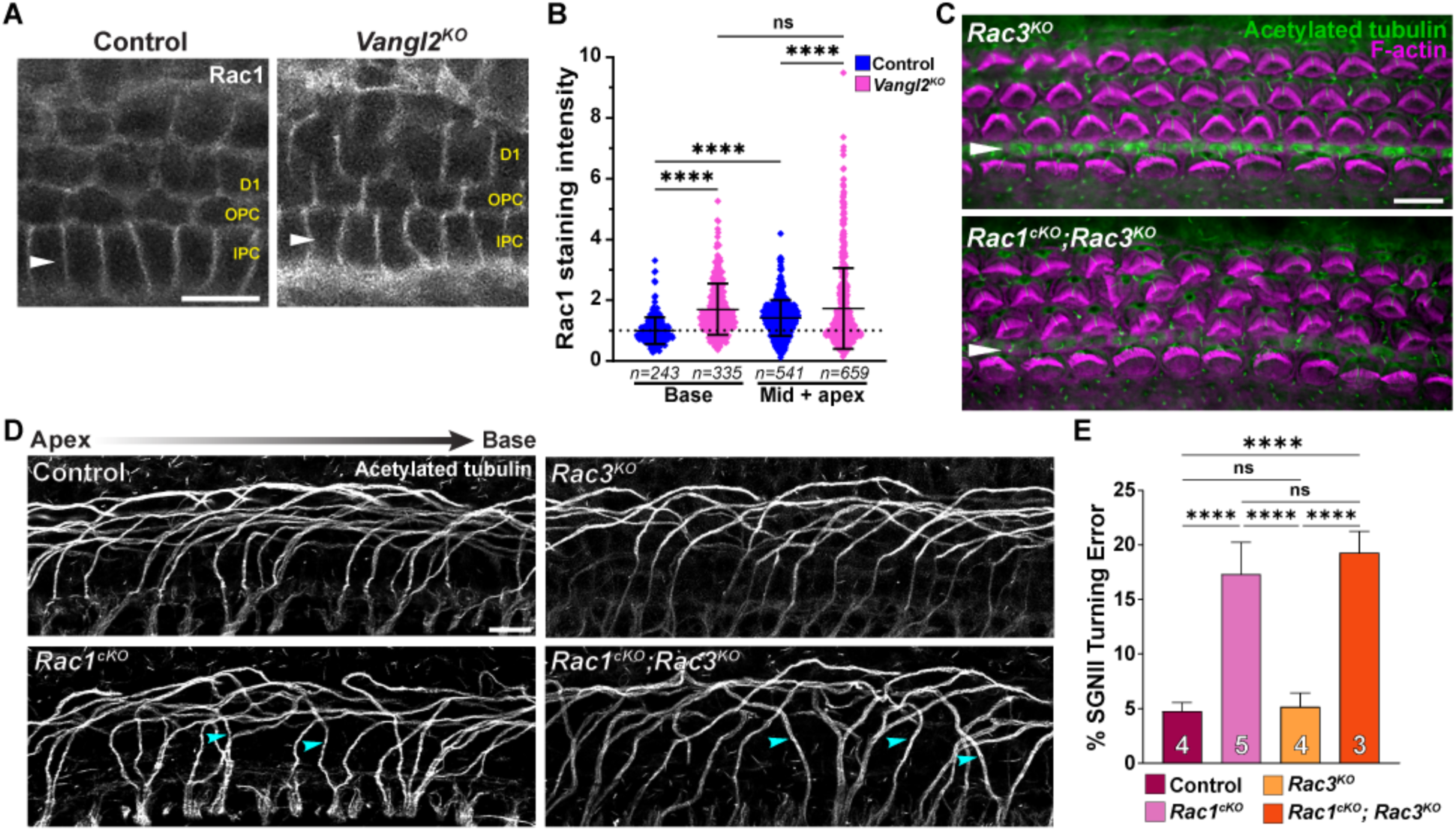
Rac1 is required in the cochlear epithelium for proper SGNII afferent turning. **A**, Rac1 localization at the SC-SC junctions in E18.5 control and *Vangl2^KO^* cochleae. White arrowheads mark the IPC row. **B,** Quantifications of Rac1 staining intensity across apex- and base-facing SC-SC junctions, normalized to the base of the control cochleae in each experiment. Control N = 3; *Vangl2^KO^* N = 4. Total numbers of cell junctions quantified in each group are indicated by *n*. **C,** HBs and kinocilia in *Rac3^KO^* and *Rac1^cKO^*; *Rac3^KO^*E18.5 cochlea labeled by F-actin (magenta) and acetylated tubulin (green) staining, respectively. **D,** SGNII afferents labeled by acetylated tubulin staining in E18.5 control, *Rac1^cKO^*, *Rac3^KO^*, and *Rac1^cKO^*; *Rac3^KO^* cochleae. Afferents turning errantly toward the apex are denoted by cyan arrowheads. **E,** Quantifications of SGNII afferents turning errantly toward the apex in the indicated genotypes. Number of mice represented in each column is notated. For all graphs: Mean +/- stdev, one-way ANOVA with Tukey’s Post-test, ns: not significant, ∗∗: p ≤ 0.01, ∗∗∗∗: p ≤ 0.0001. Scale bars: 10 µm.

### Rac1 but not Rac3 is required in the cochlear epithelium for SGNII afferent turning

To test this, we next determined whether Rac1 is required in the cochlear epithelium for SGNII afferent turning. We crossed a *Rac1* flox allele (Glogauer et al., 2003) with the *Emx2^Cre^* mouse line (Kimura et al., 2005) to eliminate *Rac1* specifically in the cochlear epithelium, leaving neuronal Rac1 expression intact. Rac1 and Rac3 are both expressed in the inner ear, share a 92% amino acid identity (Haataja et al., 1997), and have been shown to have redundant functions in many cases (Haataja et al., 1997; Grimsley-Myers et al., 2012). Therefore, in addition to *Rac1* conditional KO (hereby referred to as *Rac1^cKO^*) mutants, we generated both global *Rac3^KO^* and *Rac1^cKO^; Rac3^KO^* double mutant mice. Embryos were recovered at E18.5 and analyzed for HB orientation and SGNII afferent turning defects. As expected, we observed significant HB morphology and orientation defects in *Rac1^cKO^*mutants **(Fig. 2C)**, albeit milder and without an obvious decrease in cochlear duct length compared to pan-inner ear deletion of *Rac1* at an earlier stage using *Foxg1^Cre^* (Grimsley-Myers et al., 2009, 2012). We then analyzed the SGNII afferent turning phenotype. Importantly, the *Rac1^cKO^*and *Rac1^cKO^; Rac3^KO^* showed significant SGNII turning errors, similar in severity to each other but milder compared to the *Vangl2^KO^* **(Fig. 2D, E)**. The *Rac3^KO^* alone did not have any significant SGNII afferent misturning. Thus, Rac1 but not Rac3 is required in the cochlear epithelium to guide SGNII afferents.

### Expression of constitutively active Rac1 is insufficient to rescue PCP phenotypes in *Vangl2^cKO^* cochleae

To assess Rac1 activity at the SC-SC junctions, we performed IF on E18.5 cochleae using a previously characterized Rac1-GTP-specific antibody (Sipe et al., 2013). However, Rac1-GTP signals were barely detectable at SC-SC junctions in both control and *Vangl2^KO^* cochleae (data not shown). We therefore devised a genetic strategy to investigate whether Vangl2 could be guiding SGNII afferents by activating Rac1 signaling at SC-SC junctions. Specifically, we utilized a Cre-inducible, constitutively active Rac1-G12V transgene inserted at the Rosa26 locus (Srinivasan et al., 2009) to determine if constitutive activation of Rac1 could rescue the HB and SGNII afferent turning defects found in *Vangl2*-deficient cochleae. We used *Emx2^Cre^* to induce Rac1-G12V expression (hereby referred to as *R26-Rac1DA*) while simultaneously recombining *Vangl2* flox alleles (Song et al., 2010) to generate *Vangl2^cKO^*; *R26-Rac1DA/+* compound mutant embryos and analyzed HB misorientation and SGNII afferent guidance defects at E18.5. The *R26-Rac1DA/+* cochlea did not have significant HB misorientation or SGNII afferent misturning compared to the control **(Fig. 3A-D)**. Compared to *Vangl2^cKO^* cochleae, neither HB orientation nor SGNII afferent turning defects were significantly improved in *Vangl2^cKO^*; *R26-Rac1DA/+* cochleae **(Fig. 3A-D)**. Similar results were observed using *Pax2-Cre,* which drives recombination as early as E9.5 (Ohyama and Groves, 2004), resulting in Rac1-G12V expression in both the OC and the SGNs (data not shown). We confirmed the expression of HA-tagged Rac1-G12V in the OC, which was localized to the HB, HC-SC and SC-SC junctions **(Fig. 3E).** Therefore, expression of constitutively active Rac1 in the cochlear epithelium is not sufficient to rescue HB and SGNII afferent turning defects in *Vangl2-*deficient cochleae, suggesting that Vangl2 and Rac1 do not act in a simple linear pathway.

**Figure 3:**
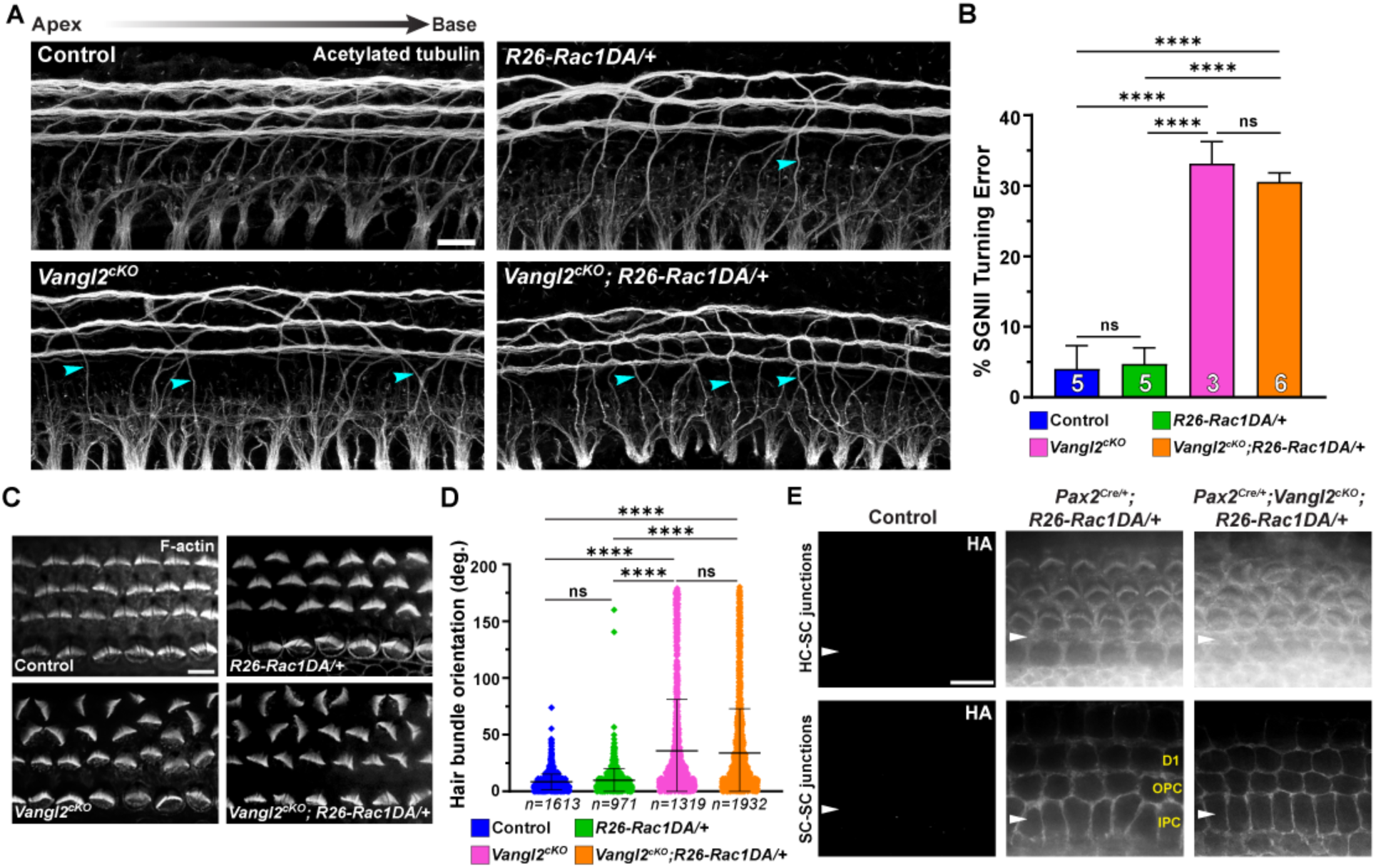
Expression of constitutively active Rac1 did not rescue *Vangl2^KO^* HB misorientation or SGNII afferent misturning. **A**, SGNII afferents labeled by acetylated tubulin staining in P0 cochleae of the indicated genotypes. Afferents turning errantly toward the apex are denoted by cyan arrowheads. **B,** Quantifications of SGNII afferents turning errantly toward the apex in the indicated genotypes. Number of samples represented in each column is noted. **C,** HBs marked by F-actin staining in the indicated genotypes. **D,** Quantifications of HB orientation in the indicated genotypes. Control N = 5; *R26-Rac1DA/+* N = 3; *Vangl2^cKO^* N = 4; *Vangl2^cKO^; R26-Rac1DA/+* N = 6. Total numbers of HCs quantified in each group are indicated by *n*. **E,** Localization of the HA-tagged Rac1DA protein in control, *Pax2-Cre; R26-Rac1DA/+*, and *Pax2-Cre; Vangl2^cKO^; R26-Rac1DA/+* P0 cochleae at the HC apical surface level (top) and SC-SC junction (bottom). White arrowheads indicate the IPC row. For all graphs: Mean +/- stdev, one-way ANOVA with Tukey’s Post-test, ns: not significant, ∗∗∗∗: p ≤ 0.0001. Scale bars: 10 µm (**A, E**), 6 µm (**C**).

### Rac1 regulates PCP protein localization at the SC-SC junctional level

To further understand the connections between Rac1 and PCP signaling in SGNII afferent guidance, we asked whether Rac1 regulates core PCP protein distribution at SC-SC junctions. In previous work, we observed a decrease in core PCP protein localization at HC-SC junctions in *Rac1-*deficient cochleae (Grimsley-Myers et al., 2009), so we analyzed distributions of representative core PCP proteins Vangl2 and Dishevelled (Dvl) at SC-SC junctions. In the control, Vangl2 localization to the SC-SC contacts had significantly higher levels at the mid and apical turns relative to the base, forming an increasing gradient along the length of the cochlea **(Fig. 4A, B)**. In *Rac1^cKO^* OC, Vangl2 localization to SC-SC junctions was decreased in the mid and apical turns but unchanged at the base relative to the control **(Fig. 4A, B)**. We also analyzed the localization of Dvl proteins, which physically interact with Vangl2 (Torban et al., 2004). In the control, Dvl3 was expressed in all SC types in the OC and localized to SC-SC contacts **(Fig. 4C)**, while Dvl2 expression in SCs was undetectable at SC-SC junctions (data not shown). In contrast to Vangl2, Dvl3 levels showed a base-apex decreasing gradient in the control. Strikingly, we found a significant decrease in Dvl3 levels along the entire length of the *Rac1^cKO^* cochleae compared to the control (**Fig. 4C, D)**. Thus, Rac1 promotes the localization of at least two core PCP proteins at SC-SC junctions. Together with the altered Rac1 localization observed in the *Vangl2^KO^*OC, these results reveal a feedback regulatory loop wherein Vangl2 regulates spatial distribution of Rac1, which in turn regulates core PCP protein localization at SC-SC contacts.

**Figure 4:**
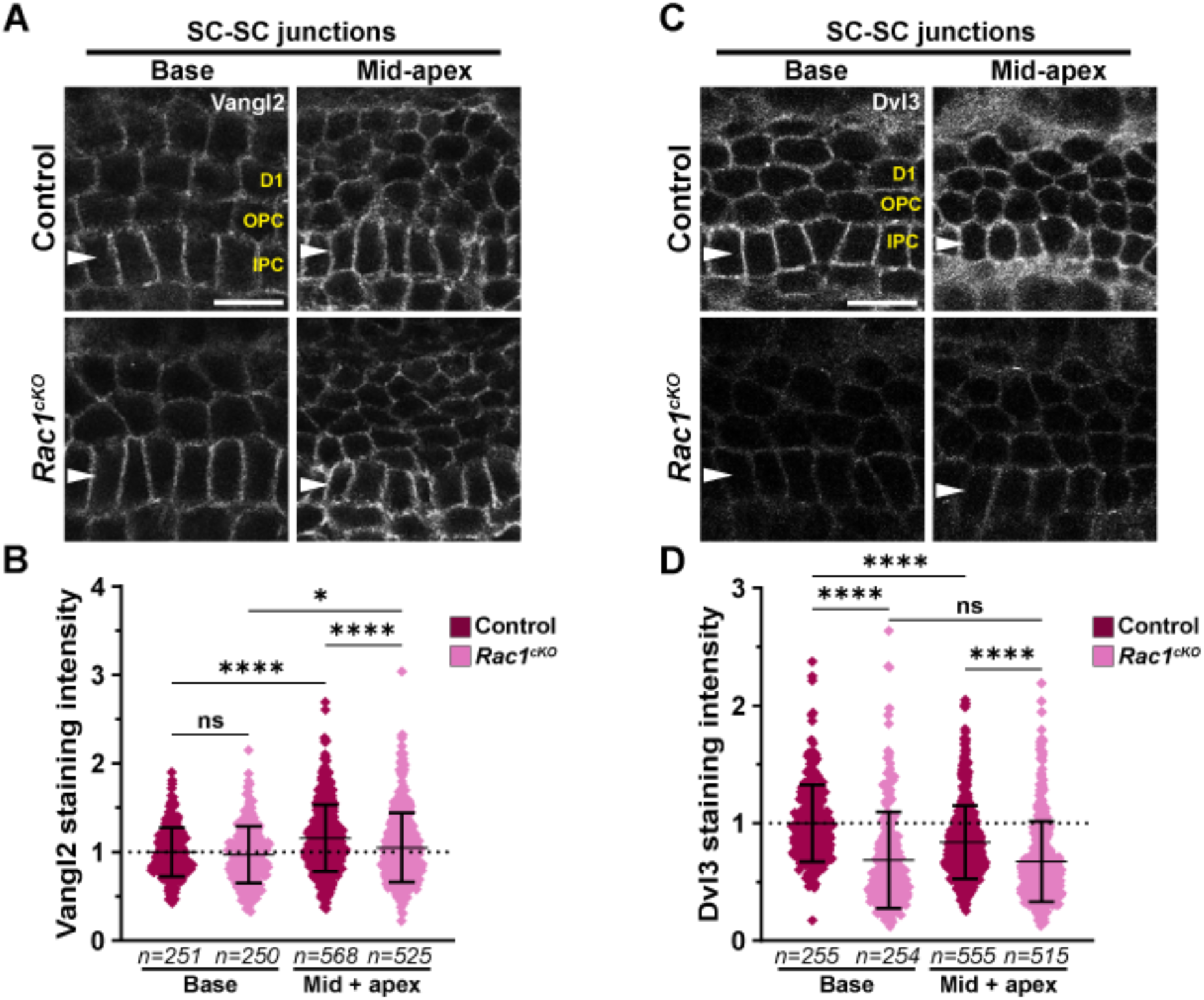
Rac1 is required in the cochlear epithelium for proper core PCP protein localization. **A, C,** Vangl2 (**A**) and Dvl3 (**C**) localization at SC-SC junctions in E17.5 control and *Rac1^cKO^* cochleae. **B, D,** Quantifications of Vangl2 (**B**, N=5 each) and Dvl3 (**D**, N=3 each) staining intensity across apex- and base-facing SC-SC junctions, normalized to the base of the control cochleae in each experiment. Total numbers of cell junctions quantified in each group are indicated by *n*. Mean +/- stdev, one-way ANOVA with Tukey’s Post-test, ns: not significant, ∗: p ≤ 0.05, ∗∗∗: p ≤ 0.001, ∗∗∗∗: p ≤ 0.0001. White arrowheads indicate the IPC row. Scale bars: 10 µm.

### Vangl2 differentially regulates Nectin3 protein localization along the apical-basal axis of cochlear SCs

The SGNII afferent turning defect in the *Rac1^cKO^* was not as severe as in the *Vangl2^KO^* (Ghimire et al., 2018), suggesting there may be other PCP effectors. Nectin3 is a good candidate because it is expressed in SCs and Nectin3 deletion causes misoriented and deformed HBs (Togashi et al., 2011; Fukuda et al., 2014). While Nectin3 has been localized to HC-SC junctions, its distribution at SC-SC junctions has not been determined. Therefore, we sought to investigate whether Nectin3 is present at SC-SC junctions during SGNII afferent turning and whether its distribution is regulated by Vangl2.

In the E18.5 control, we confirmed Nectin3 localization to HC-SC junctions and found that its levels at HC-SC junctions showed a modest base-to-apex increasing gradient **(Fig. 5A, B)**. We also detected Nectin3 at SC-SC junctions, also in a modest base-to-apex increasing gradient but at a significantly lower level than at the HC-SC junctions **(Fig. 5C)**. In the *Vangl2^KO^*, while still showing a base-to-apex increasing gradient, Nectin3 localization at the HC-SC junctions was significantly decreased in the mid and apical turns but unchanged in the basal turn relative to the control **(Fig. 5A, B)**. Interestingly, Nectin3 levels at the SC-SC junctions were significantly increased along the entire cochlear spiral **(Fig. 5C, D)**. Thus, Vangl2 regulates the spatial distribution of Nectin3 along the luminal-basal axis of SCs. Loss of Vangl2 results in decreased Nectin3 levels at HC-SC junctions but increased Nectin3 levels at SC-SC junctions; this shift in Nectin3 localization in SCs was most evident in the mid and apical turns of the cochlea. These findings suggest that Nectin3 is another likely effector of PCP signaling for SGNII afferent turning.

**Figure 5:**
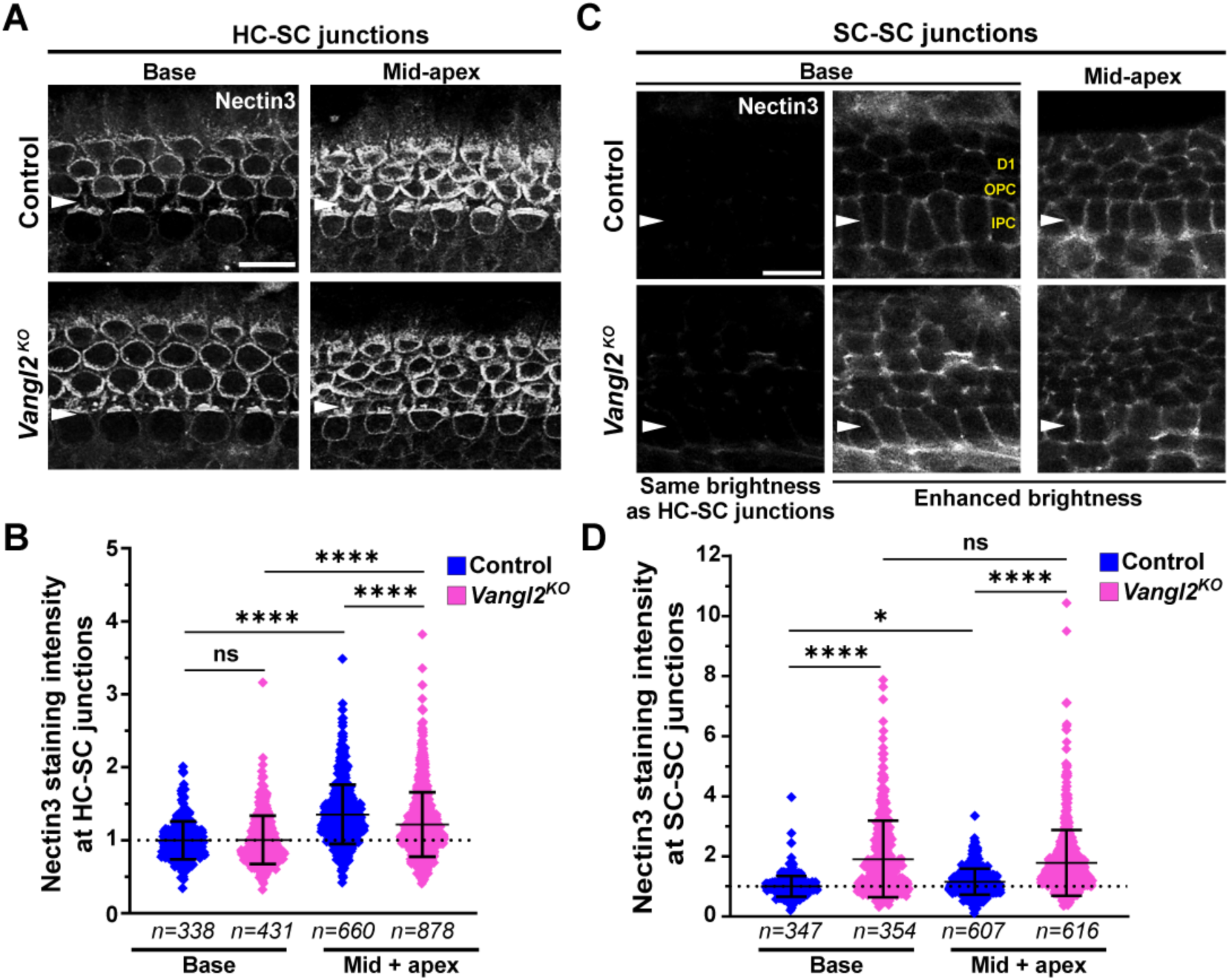
Nectin3 localization in SCs is regulated by Vangl2. **A**, Nectin3 localization to the HC-SC junctions in the basal and mid-apical regions of E18.5 control and *Vangl2^KO^*cochleae. **B,** Quantifications of Nectin3 staining intensity at the HC-SC junctions in E18.5 control and *Vangl2^KO^* cochleae normalized to the average of the littermate control basal region. For both genotypes, N = 4. **C,** Nectin3 localization at SC-SC junctions in the same samples and regions as in **A**. Panels in left column are adjusted to the same brightness as panels showing the HC-SC junctional staining, while the brightness in the middle and right columns is uniformly adjusted to optimize visibility. **D,** Quantifications of Nectin3 staining intensity across apex- and base-facing SC-SC junctions, normalized to the base of the control cochleae in each experiment. Control N = 5; *Vangl2^KO^* N = 6. For all graphs: Total numbers of cell junctions quantified in each group are indicated by *n*. Mean +/- stdev, one-way ANOVA with Tukey’s Post-test, ns: not significant, ∗∗∗: p ≤ 0.001, ∗∗∗∗: p ≤ 0.0001. White arrowheads indicate the IPC row. Scale bars: 10 µm.

Vangl2 has been shown to regulate matrix metalloproteinases (MMPs), which are endopeptidases known to degrade adhesion proteins such as Nectin3 (Williams et al., 2012; van der Kooij et al., 2014; Jessen and Jessen, 2017). Therefore, a potential mechanism by which Vangl2 regulates Nectin3 localization is through modulating MMP localization and/or activity. As Vangl2 has been reported to regulate MMP14 trafficking during zebrafish gastrulation (Williams et al., 2012), we characterized MMP14 localization in the *Vangl2^KO^*cochlea at E18.5 **(Supplementary Fig. 1)**. In both the control and *Vangl2^KO^* cochlea, MMP-14 showed a punctate staining pattern in the OHC region and localized to the apical junctions of IHCs and cells in the greater epithelial ridge (GER). Of note, no MMP14 IF signals were detected at the SC-SC junctions in either the control or *Vangl2^KO^* cochlea (data not shown). Therefore, our results do not support a role for Vangl2 in the regulation of MMP14 trafficking in the OC.

### Nectin3 is necessary for SGNII afferent turning

To investigate whether Nectin3 is necessary for SGNII afferent guidance, we generated two *Nectin3* KO alleles via CRISPR/Cas9-mediated gene editing: the *Nectin3^del10^* allele carries a 10-base-pair deletion in the beginning of exon 2 that is predicted to cause a frameshift and early STOP codon, while the *Nectin3^dE2^* allele deleted most sequences in exon 2 and also introduced an early STOP **(Supplementary Fig. 2A, B)**. After sequencing and backcrossing, we verified that both mutant lines phenocopied the reported null allele, including male sterility and microphthalmia (Inagaki et al., 2005). Moreover, both the *Nectin3^del10/del10^* and the *Nectin3^dE2/dE2^* OC displayed defective cellular mosaic in the OC and aberrant HC-HC contact sites, misoriented HBs, and mispositioned kinocilia characteristic of published *Nectin3* KO mouse lines **(Supplementary Fig. 2C)** (Togashi et al., 2011). We next assessed SGNII afferent turning in P0 control, heterozygous, and mutant OC of both *Nectin3* KO alleles **(Fig. 6A-E)**.

**Figure 6:**
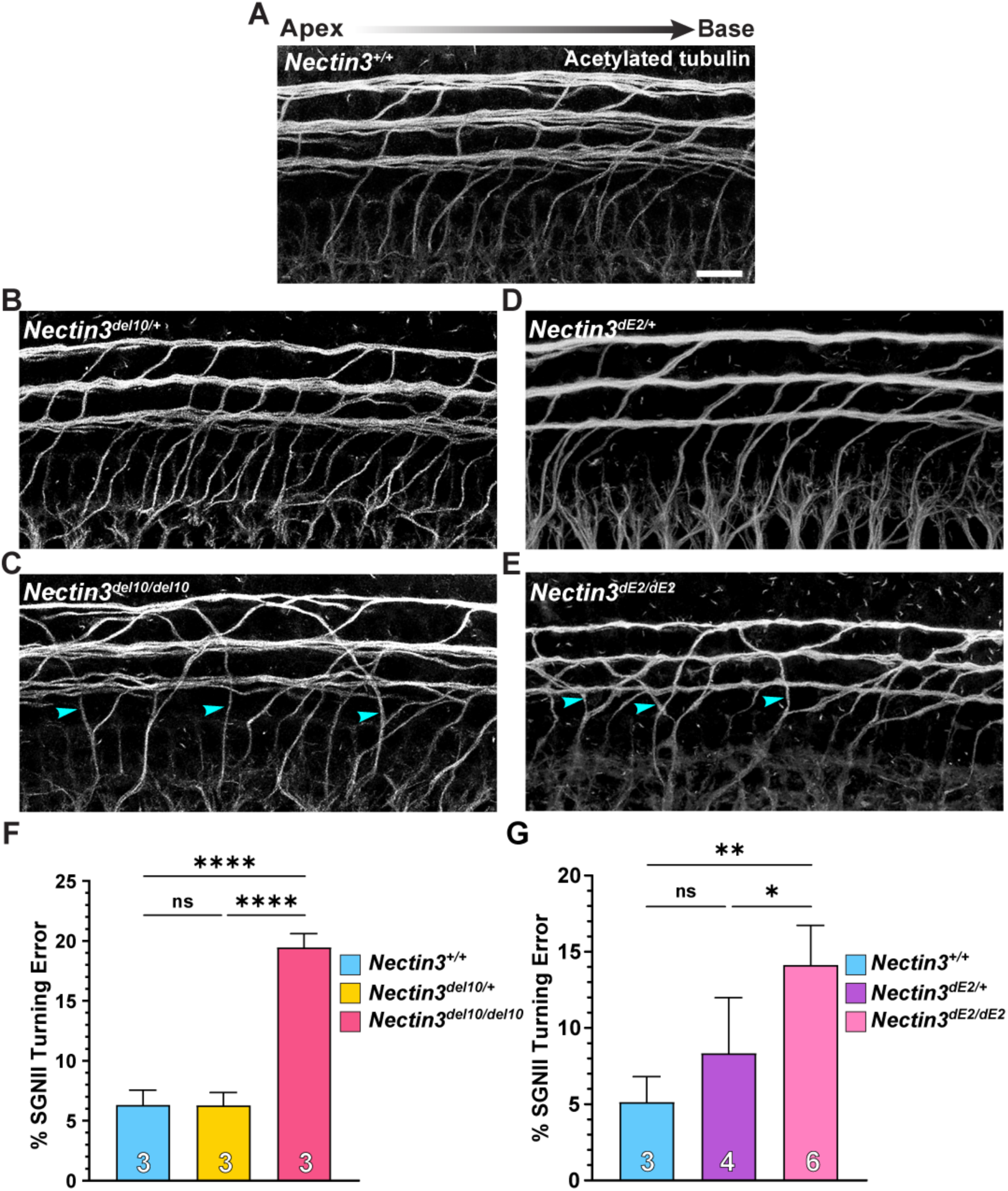
Nectin3 is necessary for proper SGNII afferent turning. **A-E,** SGNII afferents labeled by acetylated tubulin staining in P0-P1 cochleae of the indicated genotypes, with errantly turning afferents denoted by cyan arrowheads. **F, G,** Quantifications of SGNII afferents errantly turning toward the apex in the indicated *Nectin3^del10^*(**F**) and *Nectin3^dE2^* genotypes (**G**). Although both alleles are predicted to be null alleles, the *Nectin3^del10/del10^* had significantly higher SGNII afferent misturning compared with the *Nectin3^dE2/dE2^* (unpaired Student’s T-test, p=0.0133, comparison not shown). For all graphs: Mean +/- stdev, One-way ANOVA with Tukey’s Post-test, ns: not significant, ∗: p ≤ 0.05; ∗∗: p ≤ 0.01, ∗∗∗∗: p ≤ 0.0001. Scale bar: 10 µm.

Remarkably, both mutants showed significant SGNII afferent misturning. Upon quantification, the severity of SGNII afferent misturning in both the *Nectin3^del10/del10^* and *Nectin3^dE2/dE2^* cochleae was similar to that of *Rac1^cKO^* cochleae but milder than in *Vangl2^KO^*cochleae **(Fig. 6F, G)**. These results indicate that Nectin3 is necessary for proper SGNII afferent turning.

### Nectin3 and Rac1 likely act in the same genetic pathway to regulate SGNII afferent turning

Because *Rac1^cKO^* and *Nectin3^del10/del10^*, hereafter *Nectin3^KO^*, cochleae had similar SGNII afferent turning defects compared to each other but milder compared to *Vangl2^KO^*, we investigated whether Nectin3 and Rac1 act in concert for SGNII afferent guidance. We generated a compound *Nectin3^KO^; Rac1^cKO^* mutant and characterized the SGNII afferent turning phenotype **(Fig 7A-C)**. Interestingly, the severity of SGNII afferent turning defects in the *Nectin3^KO^; Rac1^cKO^* was similar to that found in either single mutant **(Fig. 7D)**. Likewise, the aberrant HC-HC contacts and kinocilium positioning defects were not worsened in the double mutant compared to *Nectin3^KO^* **(Supplementary Fig. 3A-C)**. On the other hand, there was a modest but significant increase in HB misorientation in *Nectin3^KO^; Rac1^cKO^* cochleae compared to each of the single mutants **(Supplementary Fig. 3D)**. These results are consistent with Rac1 and Nectin3 acting in the same pathway to regulate SGNII afferent turning.

**Figure 7:**
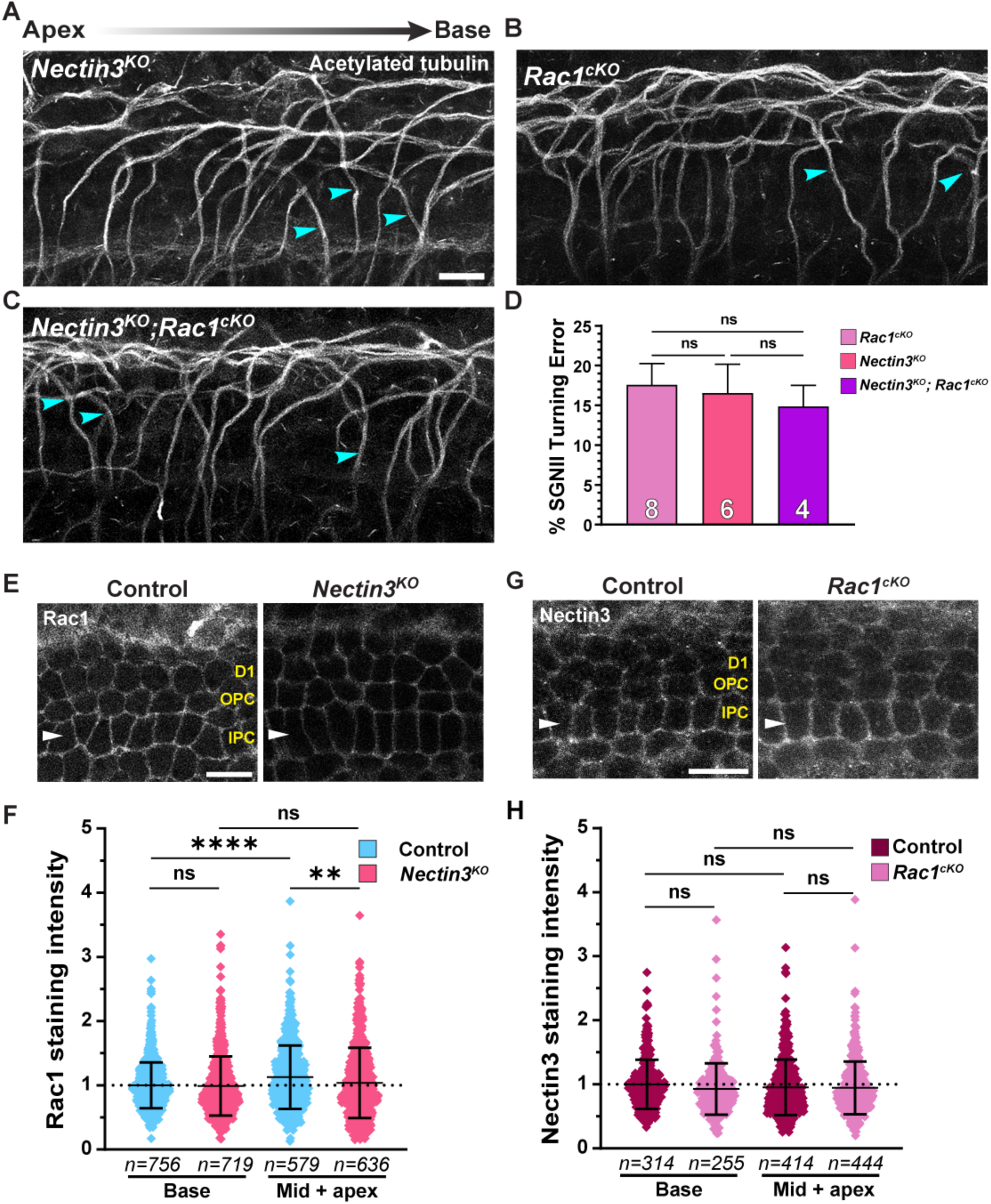
Nectin3 and Rac1 likely act in the same genetic pathway to regulate SGNII afferent turning. **A-C,** SGNII afferents labeled by acetylated tubulin staining in E18.5-P0 *Nectin3^KO^* (**A**), *Rac1^cKO^*(**B**), and *Nectin3^KO^; Rac1^cKO^* (**C**) cochleae, with examples of errantly turning afferents denoted by cyan arrowheads. **D,** Quantification of SGNII afferents errantly turning toward the apex in the indicated genotypes. **E,** Rac1 localization at the SC-SC junctions of control and *Nectin3^KO^* E18.5 cochleae. White arrowheads indicate the IPC row. **F,** Quantification of Rac1 staining intensity across apex- and base-facing SC-SC junctions, normalized to the base of the control cochlea in each experiment. N = 5 for both genotypes. **G,** Nectin3 localization at the SC-SC junctions of Control and *Rac1^cKO^* E18.5 cochleae. White arrowheads indicate the IPC row. **H,** Quantification of Nectin3 staining intensity across apex- and base-facing SC-SC junctions, normalized to the base of the control cochlea in each experiment. N = 3 for both genotypes. For all graphs: Total numbers of cell junctions quantified in each group are indicated by *n*. Mean +/- stdev, one-way ANOVA with Tukey’s Post-test, ns: not significant, ∗: p ≤ 0.05, ∗∗∗: p ≤ 0.001, ∗∗∗∗: p ≤ 0.0001. Scale bar: 10 µm.

Nectin3-mediated adhesion can activate Rac1 in cultured cells (Ogita and Takai, 2006). To test whether Rac1 acts downstream of Nectin3 in the OC, we first examined Rac1 localization at the SC-SC junctions of *Nectin3^KO^* cochleae. Rac1 levels were unchanged at the base but significantly decreased in the mid and apical turns compared with the control, thereby flattening the base-to-apex increasing gradient present in the control **(Fig. 7E, F)**. Conversely, Nectin3 localization at the SC-SC junctions in *Rac1^cKO^* cochleae was similar to the control **(Fig. 7G, H)**. Together, these data suggest that Rac1 may be acting downstream of Nectin3 to regulate SGNII afferent guidance.

## Discussion

In this study, we have identified the small GTPase Rac1 and the IgCAM Nectin3 as novel regulators of PCP-directed SGNII peripheral afferent guidance. We show that Rac1 and Nectin3 are localized to the junctions between SCs, which serve as intermediate targets of SGNII peripheral afferents *en route* to OHCs, and both are required for SGNII afferent turning. The epithelium-specific *Rac1^cKO^* revealed a non-autonomous role of Rac1 in the environment to guide SGNII afferent turning, thus expanding the mechanisms of action of Rac GTPases beyond their well-established cell-autonomous functions in axon guidance (Ng et al., 2002; Tahirovic et al., 2010; Hua et al., 2015). Although the site of action of Nectin3 could not be unambiguously determined using *Nectin3* global KO mutants, Nectin3 protein was not detected on SGNII peripheral processes during the time they turn towards the cochlear base (data not shown). Thus, our data favor a non-autonomous role of Nectin3 in cochlear SCs for guiding SGNII afferents.

Both Vangl2 and its ligand Fzd3/6 act non-cell-autonomously in the cochlear epithelium to guide SGNII afferents (Ghimire et al., 2018; Ghimire and Deans, 2019), suggesting that the core PCP proteins themselves are unlikely to serve as axon guidance cues for SGNII afferents. Our study suggests that PCP signaling modulates the mechanical and adhesive properties of axon guidance substrates via regulation of Rac1 and Nectin3 at SC-SC junctions (**Fig. 8A**). Of note, this mechanism is distinct from PCP-directed anterior turning of spinal cord commissural axons, where PCP proteins expressed in the growth cones guide commissural axons in response to an anterior-posterior Wnt gradient (Lyuksyutova et al., 2003; Shafer et al., 2011).

**Figure 8:**
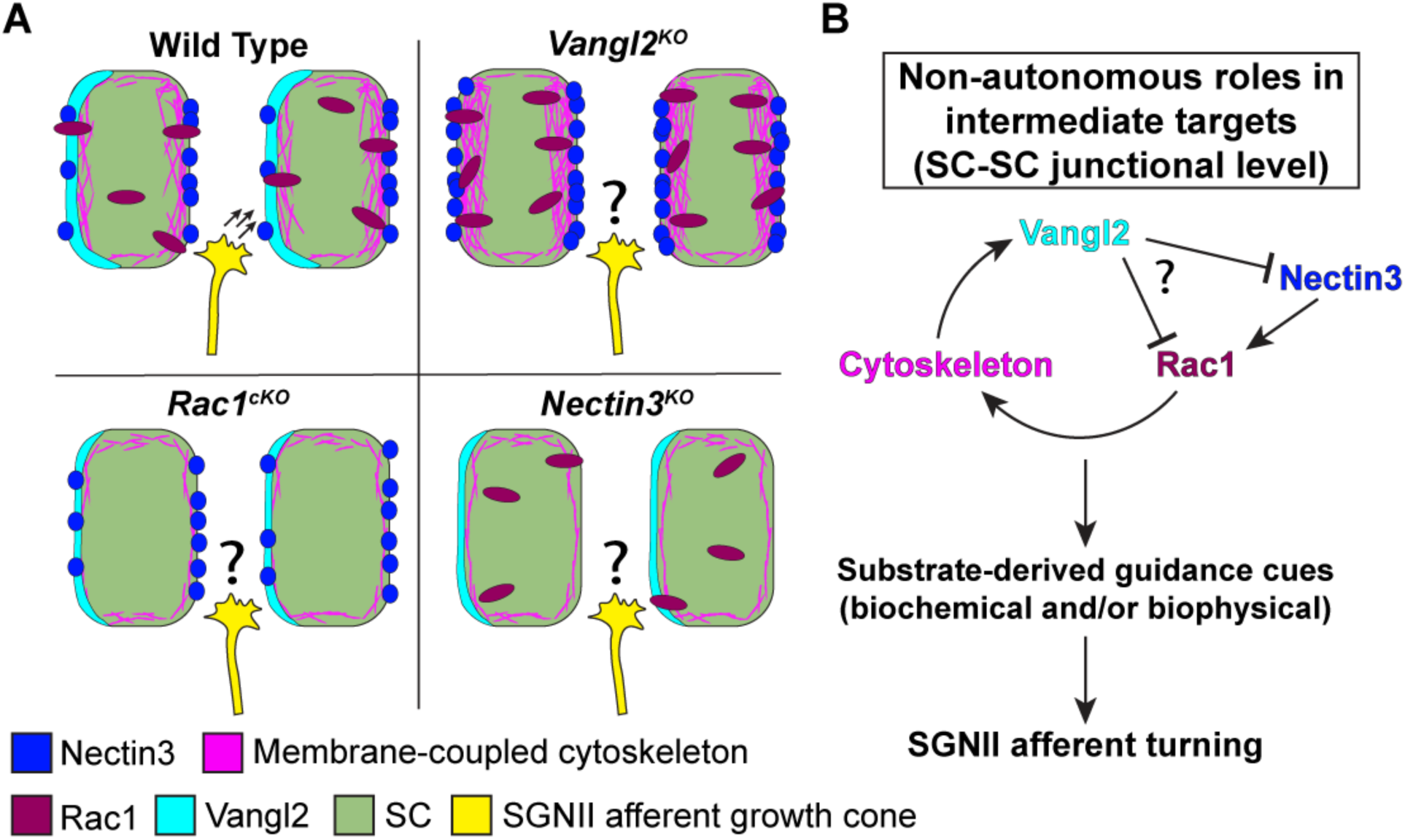
Proposed model of SC-derived SGNII afferent guidance cues. **A**, A proposed “Goldilocks” model for optimal substrate properties required for SGNII afferent turning. Both increased presence of Rac1 and Nectin3, as occurred in *Vangl2^KO^*, and their absence, impairs substrate-derived axon guidance cues for SGNII afferent turning. **B,** Genetic evidence supports a regulatory loop for protein localization of Vangl2, Nectin3, and Rac1 at SC-SC junctions to guide SGNII afferents. Rac1 also regulates Dvl3 localization of SC-SC junctions.

As a master regulator of PCP establishment (Dreyer et al., 2022), Vangl2 has been shown to regulate the endocytosis of several integral membrane proteins, such as nephrin (Babayeva et al., 2013) and Fzd3 (Shafer et al., 2011). Intriguingly, we found that Vangl2 regulates the subcellular distribution of Nectin3 along the apical-basal axis of SCs; Vangl2 loss results in decreased Nectin3 levels at HC-SC junctions but increased Nectin3 at SC-SC junctions, suggesting a spatial shift via an indirect regulatory mechanism. Vangl2 has been shown to regulate MMP trafficking and activity, which are known to degrade extracellular matrix and adhesion proteins (Williams et al., 2012; van der Kooij et al., 2014; Jessen and Jessen, 2017). While we did not find evidence for Vangl2 regulation of MMP14 trafficking in the cochlea, other MMPs may be involved. Alternatively, Vangl2 has been shown to promote coupling of cell adhesion molecules to the actin cytoskeleton (Dos-Santos Carvalho et al., 2020), and may likewise promote Nectin3-F-actin coupling at the SC-SC junctions. Because there is evidence that Vangl2 is enriched on the side of SCs facing the cochlear apex (**Fig. 1B**) (Ghimire et al., 2018), it may promote Nectin3 asymmetry on SC-SC junctions to guide SGNII afferents. Our Nectin3 immunolocalization results are not of sufficient resolution to discern whether there is Nectin3 asymmetry at the SC-SC junctions. In the future, it would be informative to determine precise Nectin3 localization by mosaic labeling.

Similarly, our previous and current findings show that Vangl2 spatially regulates Rac1 subcellular localization and activity in the OC (Grimsley-Myers et al., 2009). In turn, Rac1 promotes Vangl2 and Dvl3 localization at SC-SC junctions, forming a feedback regulatory loop (**Figure 8B**). Uniform expression of R26-*Rac1DA* was not able to restore axon guidance cues in the absence of Vangl2; Rac1DA may be insufficient to recapitulate features of the said feedback regulatory loop, or obscure any polarized distribution of Nectin3 and Rac1 activity necessary for guiding SGNII afferents. Considering increased Rac1 localization to SC-SC junctions in the *Vangl2^cKO^*OC, it is also plausible that Rac1 activity is increased at SC-SC junctions in the *Vangl2^cKO^* compared to the control. In this scenario, further activation of Rac1 in *Vangl2^cKO^* mutants would not be expected to rescue SGNII afferent turning defects.

It is worth noting that there appears to be a subcellular gradient of Vangl2, Nectin3, and active Rac1 along the apical-basal axis of SCs. Both Vangl2, Nectin3, and Rac1-GTP levels were significantly lower at SC-SC junctions compared to HC-SC junctions, suggesting weaker cell adhesions at the basal SC-SC junctions compared to the apical HC-SC junctions. Fine-tuning of SC-SC adhesion and SC-growth-cone interactions would allow optimal axon navigation. As growth cones navigate their environment, they form transient adhesions with surrounding cells to generate the traction force necessary to complete their trajectory (Vitriol and Zheng, 2012).

The substrate with which the growth cone interacts must be sufficiently adhesive for the growth cone to get traction, but not so adhesive that it impedes growth cone advance (Kerstein et al., 2015). Consistent with this idea, we observed an increase of both Rac1 and Nectin3 at SC-SC junctions of *Vangl2^KO^* cochleae compared to the control, suggesting that too much adhesion between SCs or SCs and SGNII growth cones may temporarily stall SGNII growth cones leading to turning errors (**Figure 8A**). Future experiments to overexpress Nectin3 in SCs will distinguish whether an increase of Nectin3 at SC-SC junctions itself causes SGNII afferent misturning.

Nectin3 may play both adhesion and signaling roles in SCs to guide SGNII afferents. Nectin3 homophilic adhesion between SCs is expected to be of lower affinity than the Nectin1-Nectin3 heterophilic adhesions at HC-SC junctions, thus allowing SGNII afferents to travel through unimpeded. Nectin3 in SCs may also interact with Nectin3 ligands expressed in SGNII afferents to advance SGNII growth cones. There is a precedent for this scenario in commissural axon turning in the spiral cord. Nectin3 and Nectin-like molecules expressed in the floor plate cells interact with their respective Nectin ligands in the post-crossing commissural axons to direct their anterior turning behavior (Okabe et al., 2004; Niederkofler et al., 2010). However, to our knowledge, our study is the first to uncover a functional link between Nectins and non-autonomous roles of PCP signaling in axon guidance.

We present evidence that Nectin3 and Rac1 act in the same genetic pathway in SCs for SGNII axon guidance. First, we observed similar changes in their localization at SC-SC junctions in the *Vangl2^KO^* cochlea. Second, the severity of SGNII afferent misturning rates was similar in the *Rac1^cKO^*and *Nectin3* KO mutants. Third, we did not observe additive defects in SGNII afferent turning in the *Nectin3^KO^; Rac1^cKO^* cochlea. Interestingly, the HB misorientation in the *Nectin3^KO^; Rac1^cKO^*was worse than in either mutant alone, suggesting that Rac1 likely has Nectin3- independent functions in HCs to regulate HB orientation, whereas Nectin3 and Rac1 act in the same pathway in SCs to direct SGNII afferent turning. Thus, our results argue that these two PCP-mediated functions are spatially segregated and differentially regulated at the HC-SC and SC-SC junctions, and SGNII afferent misturning does not always mirror the severity of HB misorientation. Nectin1/3 *trans* interactions have been shown to induce Rac activity in fibroblasts (Kawakatsu et al., 2002; Honda et al., 2003; Ogita and Takai, 2006), so Nectin3 may regulate Rac1 activity at both HC-SC and SC-SC junctions. Consistent with this, we show that Rac1 levels were decreased at the SC-SC junctions in *Nectin3* KO mutants but not *vice versa*. Taken together, these data support a Nectin3-Rac1 signaling axis in SCs that influences PCP-mediated guidance cues for SGNII peripheral axon turning (**Figure 8B**).

## Materials and Methods

### Mice

Animal care and use were performed in compliance with National Institutes of Health guidelines and the Animal Care and Use Committee at the University of Virginia. *Vangl2* KO and flox (Song et al., 2010), *Emx2^Cre^* (Kimura et al., 2005), *Pax2-Cre* (Ohyama and Groves, 2004), *Rac1* flox (Glogauer et al., 2003), *Rac3* KO (Cho et al., 2005), and *R26-LSL-Rac1DA* (Srinivasan et al., 2009) lines were described previously. All *KO* or *cKO* alleles were determined to be phenotypically recessive, so controls include both wild-type and heterozygous animals. For Cre-driven crosses, controls include both Cre-positive and Cre-negative animals. For timed pregnancies, the morning of the plug was designated as E0.5, and the day of birth postnatal day 0 (P0).

### Generation of Nectin3 KO alleles

*Nectin3* KO alleles were generated using CRISPR-assisted genome editing in mouse zygotes performed by the Genetically Engineered Murine Model Core Facility. The guide RNA sequence around the start of exon 2 was designed using CRISPR/Cas9 target online predictor (CCTop). crRNA, tracrRNA, ssODN, and Cas9 protein were purchased from IDT (Coralville, Iowa). crRNA and tracrRNA were diluted to 100uM in RNase-free microinjection buffer (10mM of Tris-HCl, pH 7.4, 0.25mM of EDTA). Ribonucleic protein (RNP) complex was formed by mixing and incubating Cas9 at 0.2 mg/ml with the crRNA/tracrRNA at 2mM in RNase-free microinjection buffer at 37°C for 10 minutes. ssODN was added at 0.2 mg/ml. The RNP complex with ssODN was co-delivered into the fertilized eggs from B6SJL mice by electroporation with a NEPA21 super electroporator (Nepa Gene Co., Ltd. Chiba, Japan). Following overnight culturing, two-cell zygotes were implanted into the oviducts of pseudopregnant foster mothers of ICR strain. Pups born from the foster mothers were screened using tail snip DNA by PCR genotyping followed by Sanger sequencing. Founders were backcrossed with wild-type B6J mice 3 times and offspring were used for intercrosses. Sequences of guide RNA, ssODN, and genotyping and sequencing primers are available upon request.

### Immunohistochemistry

P0 or E18.5 mouse skulls or temporal bones were dissected and fixed with 4% paraformaldehyde (PFA) for 40 minutes at room temperature (RT) or overnight at 4 degrees Celsius, or in 10% trichloroacetic acid (TCA) for 1 hour on ice. After fixation, skulls or temporal bones were washed three times in phosphate buffered saline (PBS). Cochleae were dissected and blocked in PBS with 0.1% Triton X-100, 5% heat-inactivated horse serum, and 0.02% NaN_3_ for 1 hour at RT, then incubated with primary antibodies for 18-48 hours at 4 degrees Celsius. Samples were then washed three times in PBS/0.1% Triton X-100 (PBST) before incubation with secondary antibodies, phalloidin, and Hoechst for 2 hours at RT. PFA-fixed samples were washed in PBST, post-fixed for 15 minutes at RT in 4% paraformaldehyde, then washed twice more in PBST. TCA-fixed samples were washed three times in PBST. Stained samples were flat mounted in Fluoromount-G.

The antibodies used for immunohistochemistry are listed below:

| Antibody | Concentration | Vendor | Catalog# |
| --- | --- | --- | --- |
| Alexa-conjugated phalloidin | 1:200 | Thermo Fisher | A12379, A12380, A12381, A22287 |
| Alexa-conjugated secondary antibodies | 1:500 | Thermo Fisher | A11036, A11029, A11057, A11077 |
| anti-acetylated tubulin | 1:400 | Sigma | T6793 |
| anti-Dvl2 | 1:100 (TCA) | Proteintech | 12037-1-AP |
| anti-Dvl3 | 1:200 | Proteintech | 13444-1-AP |
| anti- MMP-14 | 1:100 | Sigma | ZRB1760 |
| anti-Nectin3 | 1:100 | Sigma | U3254 |
| anti-Rac1 | 1:400 | BD Biosciences | 610651 |
| anti-Vangl2 | 1:200 | Millipore | MABN750 |

### Microscopy and image analysis

Control and mutant cochleae were imaged under identical conditions in each experiment. HBs were imaged with a Deltavision deconvolution microscope using a 60X/1.35 NA oil-immersion objective controlled via SoftWoRx software (Applied Precision). Protein localization at the SC-SC junctional level was imaged using a Leica SP8 confocal microscope built off a DMi8 confocal, using a 63X/1.4 NA oil-immersion objective controlled by LAS X software. Multiple images were taken at the basal (10% total length), mid (50% total length), and apical turns (80% total length) of the cochlear duct to capture the phenotype through the entire length of the cochlea. Images were processed using Fiji (National Institutes of Health) and Adobe Photoshop. Brightness and levels for representative image panels of HBs, F-actin at HC-SC junctions, and SGNII afferents were adjusted individually for optimum visibility. Brightness and levels for representative images of Nectin3, Rac1, Vangl2, HA, MMP14, and Dvl3 protein localization were adjusted identically.

### HB orientation quantification

The HBs were labeled with phalloidin, and the angle of rotation from the horizontal was calculated with the angle measuring function in Fiji. A HB pointing to the lateral edge of the cochlear duct has an orientation of 0 degrees, while a HB pointing parallel to the rows of hair cells in either direction would have an orientation of 90 degrees. For HB orientation, kinocilium index, and aberrant HC-HC contact quantifications, only the base and mid turns were used, as the HCs in the *Nectin3^KO^*apex were too immature for proper quantification.

### Kinocilium index quantification

Kinocilium positioning within the HB was assessed by measuring the kincocilium index, as previously described (Malt et al., 2019). Briefly, the kinocilia were labeled with acetylated tubulin, and the HBs with phalloidin. Lines were drawn and measured in Fiji from the base of the kinocilium to each end of the HB (x and y). The kinocilium index is calculated as x/y or y/x, with the largest value serving as the numerator.

### Aberrant HC-HC contacts quantification

The percentage of HCs with aberrant HC-HC contacts was counted via visualization of phalloidin at the HC-SC contact level. In cases of HC-HC contact, the rings of F-actin marking the HCs touch, usually marked by enriched F-actin.

### SGNII afferent turning quantification

SGNII afferents were labeled with anti-acetylated tubulin antibodies and traced through the Z-stack images along the length of the cochlea in Fiji. The total number of SGNII afferents that completed a turn, and the number of SGNII afferents that turned incorrectly in each image were counted. Some SGNII afferents made multiple turns, initially towards the apex and then ultimately towards the base. Such cases were not counted as misturning. The turning error was calculated as the percentage of the misturning SGNII afferents.

### Protein localization quantification

Nectin3, Rac1, Vangl2, and Dvl3 protein localization at SC-SC junctions was quantified with Fiji line scans. 1-3 µm scans were collected across the apex- and base-facing junctions and in the adjacent cell body. The average intensity value in each cell was subtracted from the maximum intensity value collected in its corresponding junction to obtain an adjusted junctional intensity. For each experiment, every value is divided by the average of the adjusted measurements from the base of the control. A value of 1 would indicate the junction has the same value as the control average, while a higher or lower value would indicate increased or decreased intensity, respectively, compared to its respective control. Nectin3 protein localization at the apical surface was also quantified with Fiji line scans across the OHC-SC junctions and in adjacent HC bodies. The average intensity value from the adjacent HC body was subtracted from the maximum intensity value across the OHC-SC junction. This adjusted junctional intensity value was normalized to the base of the control, as in the SC-SC junctional measurements.

### Statistics

Statistical analysis of cochleae from at least three mice from three different litters was performed using GraphPad Prism. Data were analyzed using a one-way analysis of variance (ANOVA) with a *post hoc* Tukey’s test, or an unpaired, two-tailed T-test. *p*-Values for statistical significance in both tests are defined as: ns = p > 0.05; ∗ = p ≤ 0.05; ∗∗ = p ≤ 0.01; ∗∗∗ = p ≤ 0.001, and ∗∗∗∗ = p ≤ 0.0001. Data were presented as mean ± standard deviation. Significance tests on SGNII afferent turning and aberrant HC-HC contact data presented in bar graphs were conducted on total values from each mouse. Significance tests on data presented in violin plots were conducted on individual cell measurements from all mice.

## Acknowledgments

We thank Wenxia Li for technical assistance, Dr. Wenhao Xu and the Genetically Engineered Murine Model Core Facility for generation of Nectin3 knockout alleles, and Dr. Yingzi Yang (Harvard School of Dental Medicine) and Dr. Karen Hirschi (University of Virginia Health System) for equipment and reagents. We thank the following organizations for funding support: NIDCD (F31-DC020667, R01-DC013773), NICHD (R01HD107872) and NIGMS (T32-GM008136), University of Virginia (Double Hoo Award, Robert R. Wagner fellowship).

## Conflict of interest statement

The authors declare no competing financial interests.

## Notes

### Competing Interest Statement

The authors have declared no competing interest.

## References

Babayeva S, Rocque B, Aoudjit L, Zilber Y, Li J, Baldwin C, Kawachi H, Takano T, Torban E (2013) Planar Cell Polarity Pathway Regulates Nephrin Endocytosis in Developing Podocytes. Journal of Biological Chemistry 288:24035–24048.

Butler MT, Wallingford JB (2017) Planar cell polarity in development and disease. Nat Rev Mol Cell Biol 18:375–388.

Cho YJ, Zhang B, Kaartinen V, Haataja L, de Curtis I, Groffen J, Heisterkamp N (2005) Generation of rac3 null mutant mice: role of Rac3 in Bcr/Abl-caused lymphoblastic leukemia. Mol Cell Biol 25:5777–5785.

Deans MR (2022) Planar cell polarity signaling guides cochlear innervation. Dev Biol 486:1–4.

Dos-Santos Carvalho S, Moreau MM, Hien YE, Garcia M, Aubailly N, Henderson DJ, Studer V, Sans N, Thoumine O, Montcouquiol M (2020) Vangl2 acts at the interface between actin and N-cadherin to modulate mammalian neuronal outgrowth Tissir F, Akhmanova A, Tissir F, eds. eLife 9:e51822.

Dreyer CA, VanderVorst K, Carraway KL (2022) Vangl as a Master Scaffold for Wnt/Planar Cell Polarity Signaling in Development and Disease. Frontiers in Cell and Developmental Biology 10 Available at: https://www.frontiersin.org/articles/10.3389/fcell.2022.887100 [Accessed July 7, 2023].

Fukuda T, Kominami K, Wang S, Togashi H, Hirata K, Mizoguchi A, Rikitake Y, Takai Y (2014) Aberrant cochlear hair cell attachments caused by Nectin-3 deficiency result in hair bundle abnormalities. Development 141:399–409.

Ghimire SR, Deans MR (2019) Frizzled3 and Frizzled6 Cooperate with Vangl2 to Direct Cochlear Innervation by Type II Spiral Ganglion Neurons. J Neurosci 39:8013–8023.

Ghimire SR, Ratzan EM, Deans MR (2018) A non-autonomous function of the core PCP protein VANGL2 directs peripheral axon turning in the developing cochlea. Development 145 Available at: https://dev.biologists.org/content/145/12/dev159012 [Accessed September 26, 2020].

Glogauer M, Marchal CC, Zhu F, Worku A, Clausen BE, Foerster I, Marks P, Downey GP, Dinauer M, Kwiatkowski DJ (2003) Rac1 Deletion in Mouse Neutrophils Has Selective Effects on Neutrophil Functions1. The Journal of Immunology 170:5652–5657.

Grimsley-Myers CM, Sipe CW, Géléoc GSG, Lu X (2009) The Small GTPase Rac1 Regulates Auditory Hair Cell Morphogenesis. J Neurosci 29:15859–15869.

Grimsley-Myers CM, Sipe CW, Wu DK, Lu X (2012) Redundant functions of Rac GTPases in inner ear morphogenesis. Developmental Biology 362:172–186.

Haataja L, Groffen J, Heisterkamp N (1997) Characterization of RAC3, a novel member of the Rho family. J Biol Chem 272:20384–20388.

Honda T, Shimizu K, Kawakatsu T, Fukuhara A, Irie K, Nakamura T, Matsuda M, Takai Y (2003) Cdc42 and Rac small G proteins activated by trans-interactions of nectins are involved in activation of c-Jun N-terminal kinase, but not in association of nectins and cadherin to form adherens junctions, in fibroblasts. Genes to Cells 8:481–491.

Hua ZL, Emiliani FE, Nathans J (2015) Rac1 plays an essential role in axon growth and guidance and in neuronal survival in the central and peripheral nervous systems. Neural Development 10:21.

Inagaki M, Irie K, Ishizaki H, Tanaka-Okamoto M, Morimoto K, Inoue E, Ohtsuka T, Miyoshi J, Takai Y (2005) Roles of cell-adhesion molecules nectin 1 and nectin 3 in ciliary body development. Development 132:1525–1537.

Jessen TN, Jessen JR (2017) VANGL2 interacts with integrin αv to regulate matrix metalloproteinase activity and cell adhesion to the extracellular matrix. Exp Cell Res 361:265–276.

Kawakatsu T, Shimizu K, Honda T, Fukuhara T, Hoshino T, Takai Y (2002) trans-Interactions of Nectins Induce Formation of Filopodia and Lamellipodia through the Respective Activation of Cdc42 and Rac Small G Proteins *. Journal of Biological Chemistry 277:50749–50755.

Kerstein PC, Nichol RH, Gomez TM (2015) Mechanochemical regulation of growth cone motility. Frontiers in Cellular Neuroscience 9:244.

Kimura J, Suda Y, Kurokawa D, Hossain ZM, Nakamura M, Takahashi M, Hara A, Aizawa S (2005) Emx2 and Pax6 Function in Cooperation with Otx2 and Otx1 to Develop Caudal Forebrain Primordium That Includes Future Archipallium. J Neurosci 25:5097–5108.

Lindqvist M, Horn Z, Bryja V, Schulte G, Papachristou P, Ajima R, Dyberg C, Arenas E, Yamaguchi TP, Lagercrantz H, Ringstedt T (2010) Vang-like protein 2 and Rac1 interact to regulate adherens junctions. J Cell Sci 123:472–483.

Lyuksyutova AI, Lu C-C, Milanesio N, King LA, Guo N, Wang Y, Nathans J, Tessier-Lavigne M, Zou Y (2003) Anterior-Posterior Guidance of Commissural Axons by Wnt-Frizzled Signaling. Science 302:1984–1988.

Malt AL, Dailey Z, Holbrook-Rasmussen J, Zheng Y, Hogan A, Du Q, Lu X (2019) Par3 is essential for the establishment of planar cell polarity of inner ear hair cells. PNAS 116:4999–5008.

Montcouquiol M, Rachel RA, Lanford PJ, Copeland NG, Jenkins NA, Kelley MW (2003) Identification of Vangl2 and Scrb1 as planar polarity genes in mammals. Nature 423:173–177.

Ng J, Nardine T, Harms M, Tzu J, Goldstein A, Sun Y, Dietzl G, Dickson BJ, Luo L (2002) Rac GTPases control axon growth, guidance and branching. Nature 416:442–447.

Niederkofler V, Baeriswyl T, Ott R, Stoeckli ET (2010) Nectin-like molecules/SynCAMs are required for post-crossing commissural axon guidance. Development 137:427–435.

Ogita H, Takai Y (2006) Activation of Rap1, Cdc42, and Rac by Nectin Adhesion System. In: Methods in Enzymology, pp 415–424 Regulators and Effectors of Small GTPases: Rho Family. Academic Press. Available at: https://www.sciencedirect.com/science/article/pii/S0076687906060307 [Accessed July 6, 2023].

Ohyama T, Groves AK (2004) Generation of Pax2-Cre mice by modification of a Pax2 bacterial artificial chromosome. genesis 38:195–199.

Okabe N, Shimizu K, Ozaki-Kuroda K, Nakanishi H, Morimoto K, Takeuchi M, Katsumaru H, Murakami F, Takai Y (2004) Contacts between the commissural axons and the floor plate cells are mediated by nectins. Developmental Biology 273:244–256.

Shafer B, Onishi K, Lo C, Colakoglu G, Zou Y (2011) Vangl2 Promotes Wnt/Planar Cell Polarity-like Signaling by Antagonizing Dvl1-Mediated Feedback Inhibition in Growth Cone Guidance. Developmental Cell 20:177–191.

Sipe CW, Liu L, Lee J, Grimsley-Myers C, Lu X (2013) Lis1 mediates planar polarity of auditory hair cells through regulation of microtubule organization. Development 140:1785–1795.

Song H, Hu J, Chen W, Elliott G, Andre P, Gao B, Yang Y (2010) Planar cell polarity breaks bilateral symmetry by controlling ciliary positioning. Nature 466:378–382.

Srinivasan L, Sasaki Y, Calado DP, Zhang B, Paik JH, DePinho RA, Kutok JL, Kearney JF, Otipoby KL, Rajewsky K (2009) PI3 Kinase Signals BCR-Dependent Mature B Cell Survival. Cell 139:573–586.

Tahirovic S, Hellal F, Neukirchen D, Hindges R, Garvalov BK, Flynn KC, Stradal TE, Chrostek-Grashoff A, Brakebusch C, Bradke F (2010) Rac1 Regulates Neuronal Polarization through the WAVE Complex. J Neurosci 30:6930–6943.

Tarchini B, Lu X (2019) New insights into regulation and function of planar polarity in the inner ear. Neurosci Lett 709:134373.

Togashi H, Kominami K, Waseda M, Komura H, Miyoshi J, Takeichi M, Takai Y (2011) Nectins Establish a Checkerboard-Like Cellular Pattern in the Auditory Epithelium. Science 333:1144–1147.

Torban E, Wang H-J, Groulx N, Gros P (2004) Independent Mutations in Mouse *Vangl2* That Cause Neural Tube Defects in *Looptail* Mice Impair Interaction with Members of the *Dishevelled* Family*. Journal of Biological Chemistry 279:52703–52713.

van der Kooij MA, Fantin M, Rejmak E, Grosse J, Zanoletti O, Fournier C, Ganguly K, Kalita K, Kaczmarek L, Sandi C (2014) Role for MMP-9 in stress-induced downregulation of nectin-3 in hippocampal CA1 and associated behavioural alterations. Nat Commun 5:4995.

Vitriol EA, Zheng JQ (2012) Growth cone travel in space and time: the cellular ensemble of cytoskeleton, adhesion, and membrane. Neuron 73:1068–1081.

Wang Y, Guo N, Nathans J (2006) The Role of Frizzled3 and Frizzled6 in Neural Tube Closure and in the Planar Polarity of Inner-Ear Sensory Hair Cells. J Neurosci 26:2147–2156.

Weisz CJC, Williams S-PG, Eckard CS, Divito CB, Ferreira DW, Fantetti KN, Dettwyler SA, Cai H-M, Rubio ME, Kandler K, Seal RP (2021) Outer Hair Cell Glutamate Signaling through Type II Spiral Ganglion Afferents Activates Neurons in the Cochlear Nucleus in Response to Nondamaging Sounds. J Neurosci 41:2930–2943.

Williams BB, Cantrell VA, Mundell NA, Bennett AC, Quick RE, Jessen JR (2012) VANGL2 regulates membrane trafficking of MMP14 to control cell polarity and migration. J Cell Sci 125:2141–2147.

Zhang KD, Coate TM (2017) Recent advances in the development and function of type II spiral ganglion neurons in the mammalian inner ear. Semin Cell Dev Biol 65:80–87.

