## Supplemental Figures for "Rac1 and Nectin3 are essential for PCP-directed axon guidance in the peripheral auditory system"

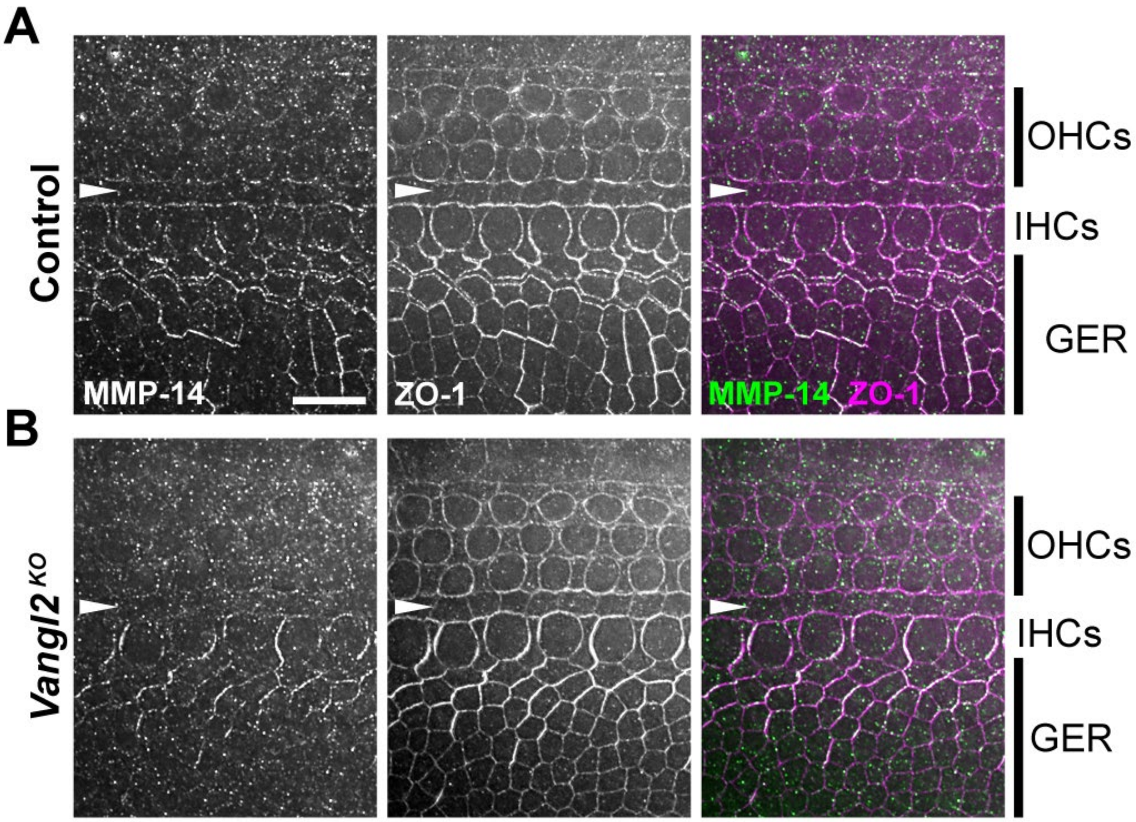


**Supplementary Figure 1: MMP-14 localization is unchanged in *Vangl2*$^{KO}$ cochleae**

**A, B,** MMP-14 localization at the HC-SC junctional level of control (**A**) and *Vangl2*$^{KO}$ (**B**). Left: MMP-14 (gray); Middle: ZO-1 (gray); Right: MMP-14 (green), ZO-1 (magenta) merge. White arrowhead indicates the IPC row, and the GER, OHC, and IHC regions are labeled on the merged panels. MMP-14 was not detected at SC-SC junctions in either genotype (not shown). Scale bar: 10 μm.

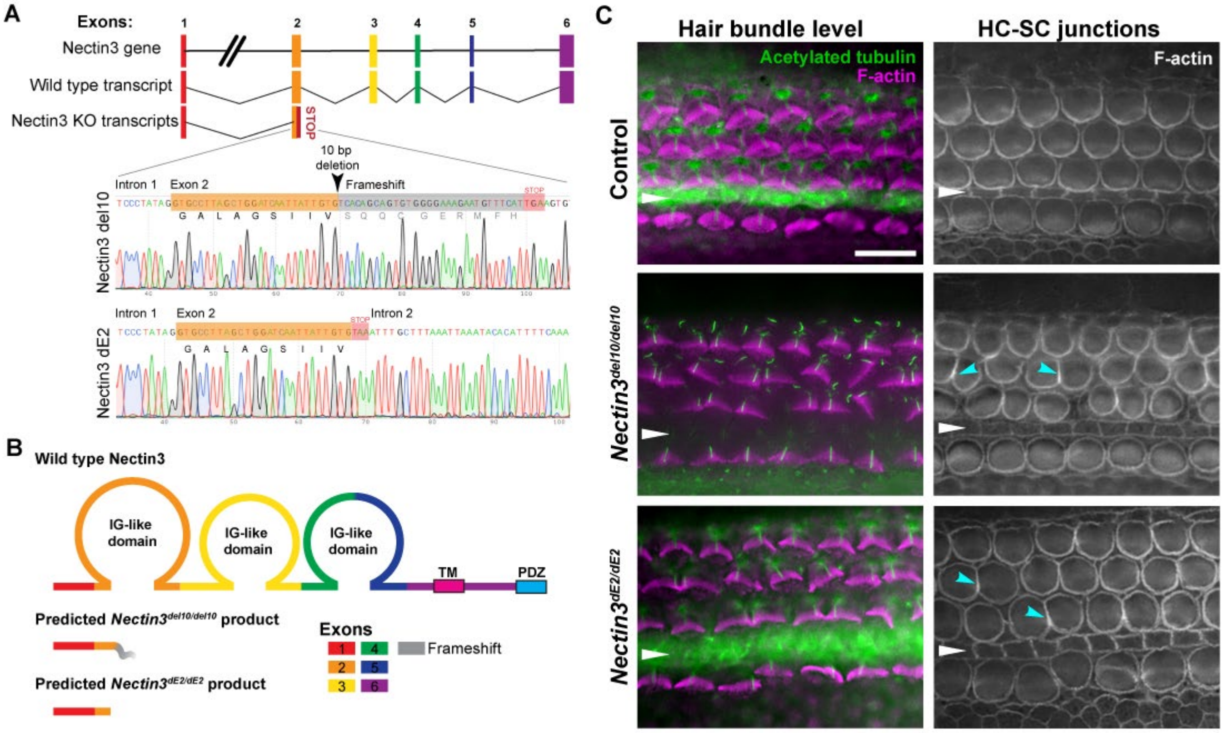


### Supplementary Figure 2: Generation of *Nectin3* knockout alleles

**A,** Schematic diagram of the Nectin3 genomic locus, predicted transcripts, and Sanger sequencing results of the *Nectin3$^{del10}$* and *Nectin3$^{dE2}$* alleles. **B,** Predicted protein product of the *Nectin3$^{del10}$* and *Nectin3$^{dE2}$* alleles. **C,** F-actin staining at the level of HBs (left, magenta) and HC-SC junctions (right, gray) of control, *Nectin3$^{del10/del10}$*, and *Nectin3$^{dE2/dE2}$* P0 cochleae. Acetylated tubulin staining (left, green) marks the kinocilium. Cyan arrowheads mark examples of enriched F-actin at aberrant HC-HC contacts. White arrowheads indicate the IPC row. Scale bar: 10 μm.

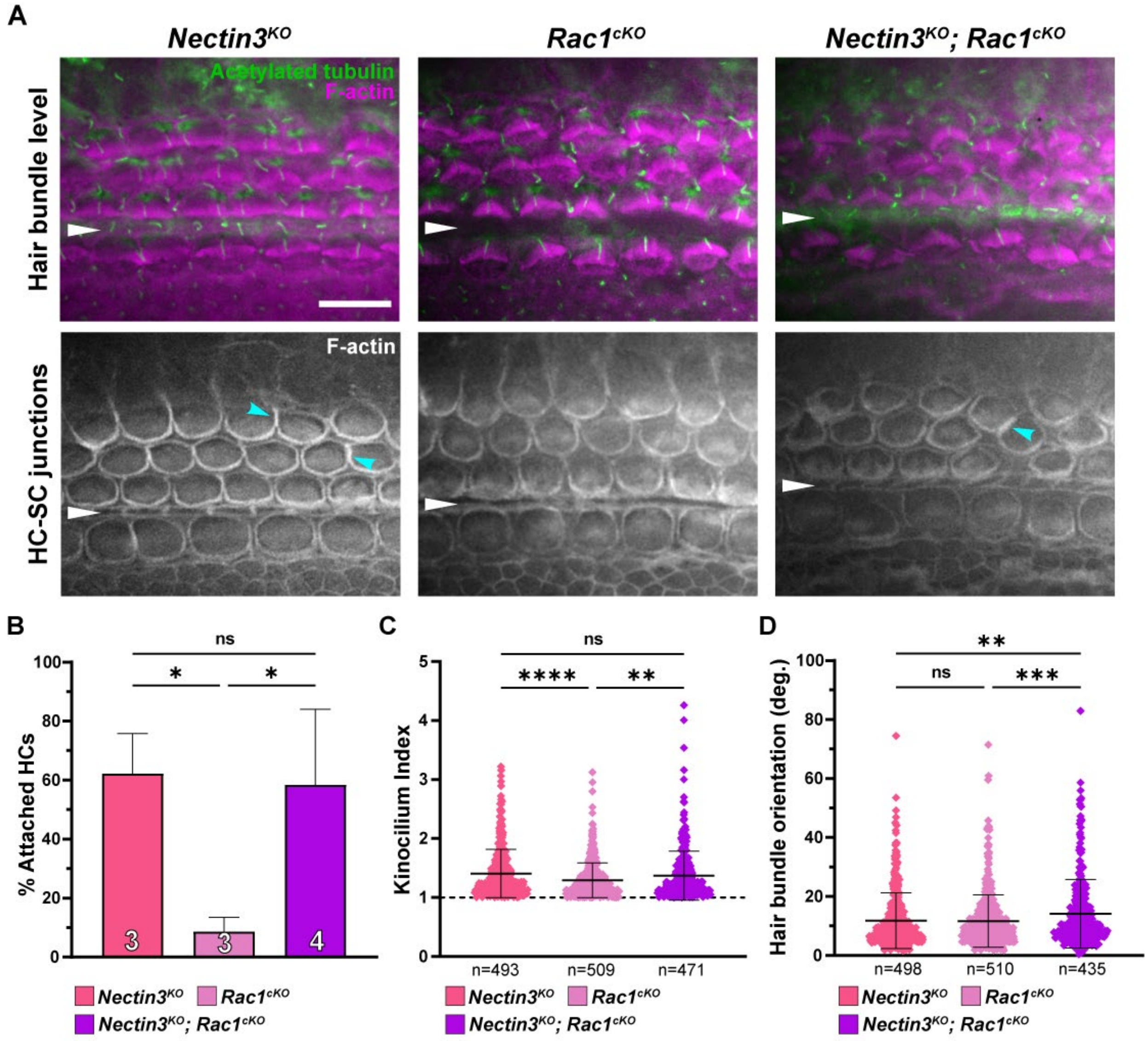


**Supplementary Figure 3: Hair cell phenotypes of Nectin3 and Rac1 compound mutants**

**A,** F-actin staining at the level of HBs (top, magenta) and HC-SC junctions (bottom, gray) of control, *Nectin3*$^{KO}$, *Rac1*$^{cKO}$*,* and *Nectin3*$^{KO}$*; Rac1*$^{cKO}$ E18-P0 cochleae. Acetylated tubulin staining (top, green) marks the kinocilium. Cyan arrowheads mark examples of enriched F-actin at aberrant HC-HC contacts. White arrowheads indicate the IPC row. Scale bar: 10 µm. **B,** Percentage of HCs aberrantly contacting other HCs in the genotypes shown in A. *Nectin3*$^{KO}$*; Rac1*$^{cKO}$ N = 4; for all other genotypes N = 3. **C, D,** Quantification of kinocilium index (**C**) and HB orientation (**D**) in the genotypes shown in A. N for all genotypes = 3. Total numbers of HCs quantified in each group are indicated by n. For all graphs: Mean +/- stdev, one-way ANOVA with Tukey's Post-test, ns: not significant, ∗: p ≤ 0.05, ∗∗: p ≤ 0.01, ∗∗∗: p ≤ 0.001, ∗∗∗∗: p ≤ 0.0001.
